# High thoughput fluorometric nucleic acid quantification using qPCR instruments

**DOI:** 10.64898/2026.08.01.742208

**Authors:** Ari Meerson

**Affiliations:** Tel Hai University of Kiryat Shmona in the Galilee, Israel; Migal Galilee Research Institute, Kiryat Shmona, Israel

## Abstract

To explore adapting qPCR systems for end-point nucleic acid quantification using dyes such as SYTO-9, we quantified serial dilutions of DNA and RNA standards in the range of 0.75 - 200 ng/µl on 384-well qPCR devices. SYTO-9 fluorescence was successfully measured using standard SYBR Green settings. Blank-subtracted relative SYTO-9 signal showed a logarithmic dependence on DNA/RNA concentration (R^2^ > 0.95). Measurements were highly stable with different incubation times, temperatures of up to 95°C, and photobleaching. The described approach is a valuable QC option for high-throughput DNA/RNA isolations and could be adapted to additional fluorometric assays beyond nucleic acids.

## Introduction

Accurate quantification of nucleic acids is a central requirement in modern molecular biology and a frequent prerequisite to downstream analyses. Among the most widely adopted approaches are fluorescence-based methods that rely on nucleic acid–binding dyes, which exhibit enhanced fluorescence upon association with double-stranded DNA (dsDNA). Since their introduction, such dyes—most notably SYBR Green I—have enabled sensitive detection of nucleic acids in solution, gels, and amplification reactions, owing to favorable photophysical properties, high selectivity for dsDNA, and strong signal enhancement upon binding (Zipper, 2003). Consequently, dye-based detection has become a cornerstone of both qualitative and quantitative nucleic acid assays.

While SYBR Green I remains the most commonly used dye, a broad range of alternatives has been developed, including the SYTO family of cyanine dyes. These dyes display diverse binding characteristics and photophysical behaviors, which can influence assay sensitivity, reaction efficiency, and reproducibility (Eischeid 2011). Among them, SYTO-9 has gained attention as a general nucleic acid stain used in applications such as flow cytometry and microscopy, as well as in PCR-based assays (Ihadjadene 2022). Importantly, SYTO-9 and SYBR Green share similar excitation and emission spectra, the latter in the green fluorescence range, enabling their detection using comparable optical configurations. This spectral overlap suggests that instrumentation optimized for SYBR Green detection may be readily adapted for use with SYTO-9 and related dyes (Jansson 2017).

Real-time quantitative PCR (qPCR) has emerged as a dominant platform for nucleic acid quantification, combining amplification and detection in a single workflow. Dye-based qPCR, particularly using SYBR Green chemistry, is widely employed due to its simplicity, cost-effectiveness, and versatility across diverse biological applications, including gene expression profiling, pathogen detection, and genotyping. In these systems, fluorescence is monitored during each amplification cycle, allowing quantification based on amplification kinetics.

Given the widespread availability of qPCR instruments in research and clinical laboratories, there is growing interest in repurposing these platforms for alternative fluorescence-based measurements. Notably, qPCR instruments are equipped with sensitive optical detection systems, precise temperature control, and multi-well plate compatibility—features that closely resemble those of conventional fluorescence plate readers. This raises the possibility of adapting qPCR systems for end-point nucleic acid quantification using DNA-binding dyes such as SYTO-9 or SYBR Green. Such an approach could offer several advantages, including improved accessibility in resource-limited settings, reduced need for specialized equipment, and streamlined workflows for high-throughput and/or automated analyses. Here, the use of qPCR instruments for end-point, fluorimetric nucleic acid quantification without amplification was evaluated.

## Materials and Methods

As input material, serial dilutions in molecular-grade water of DNA and RNA standards from DeNovix were used: 200 ng/µl DNA from the dsDNA Broad Range kit, 25 ng/µl DNA from the High Sensitivity dsDNA kit, and 100 ng/µl RNA from the RNA kit. 10 µl of the diluted DNA or RNA were mixed with 10 µl of SYTO-9 (Invitrogen #S34854) diluted to 10, 5 or 2.5 µM in molecular-grade water, mixed thoroughly and split to quadruplicates (5 µl /well) in a 384-well qPCR plate (Bio-Rad #HSP3901). The plate was covered with a Microseal ‘B’ PCR Plate Sealing Film (Bio-Rad #MSB1001B). The measurement was performed in a Bio-Rad S1000 and C1000 Touch PCR cyclers, both equipped with a 384 well block and CFX384 real-time module. The base program was 24°C for 30 sec + plate read (with standard SYBR Green filter settings). Variations of the program were also tested, e.g. repeated (3X) plate reads with averaging of RFU values per sample; pre-incubation at 95°C for 30 sec and 24°C for 30 sec before plate read, and repeated (up to 7X) reads of the same samples.

## Results

SYTO-9 fluorescence was successfully measured on our qPCR instruments, as expected, using standard SYBR Green settings.

Two representative measurement runs using serial dilutions (12.5, 25, 50, 100 and 200 ng/µl) of a 200 ng/µl DNA standard, are shown in Fig.1A,B. Using a 2.5 µM SYTO-9 working stock, RFU values between 4000 (for blank samples) and 60,000 were obtained. Blank-subtracted relative SYTO-9 fluorescence showed a logarithmic dependence on DNA concentration. When plotted on a semi-logarithmic scale, the measurements aligned well with logarithmic trendlines (R^2^ > 0.95) (Fig. 1A). Reversing the axes to facilitate the calculation of DNA concentration based on RFU, a consistent exponential dependence of the form [DNA]=Ae^0.0003F^ was observed, where [DNA] denotes DNA concentration in ng/µl, A is a coefficient experimentally established by a standard curve (in our measurements, 1<a<2), and F is the end-point, blank-subtracted RFU value. Corresponding trendlines showed R^2^ > 0.97 (Fig. 1B).

**Figure 1.**
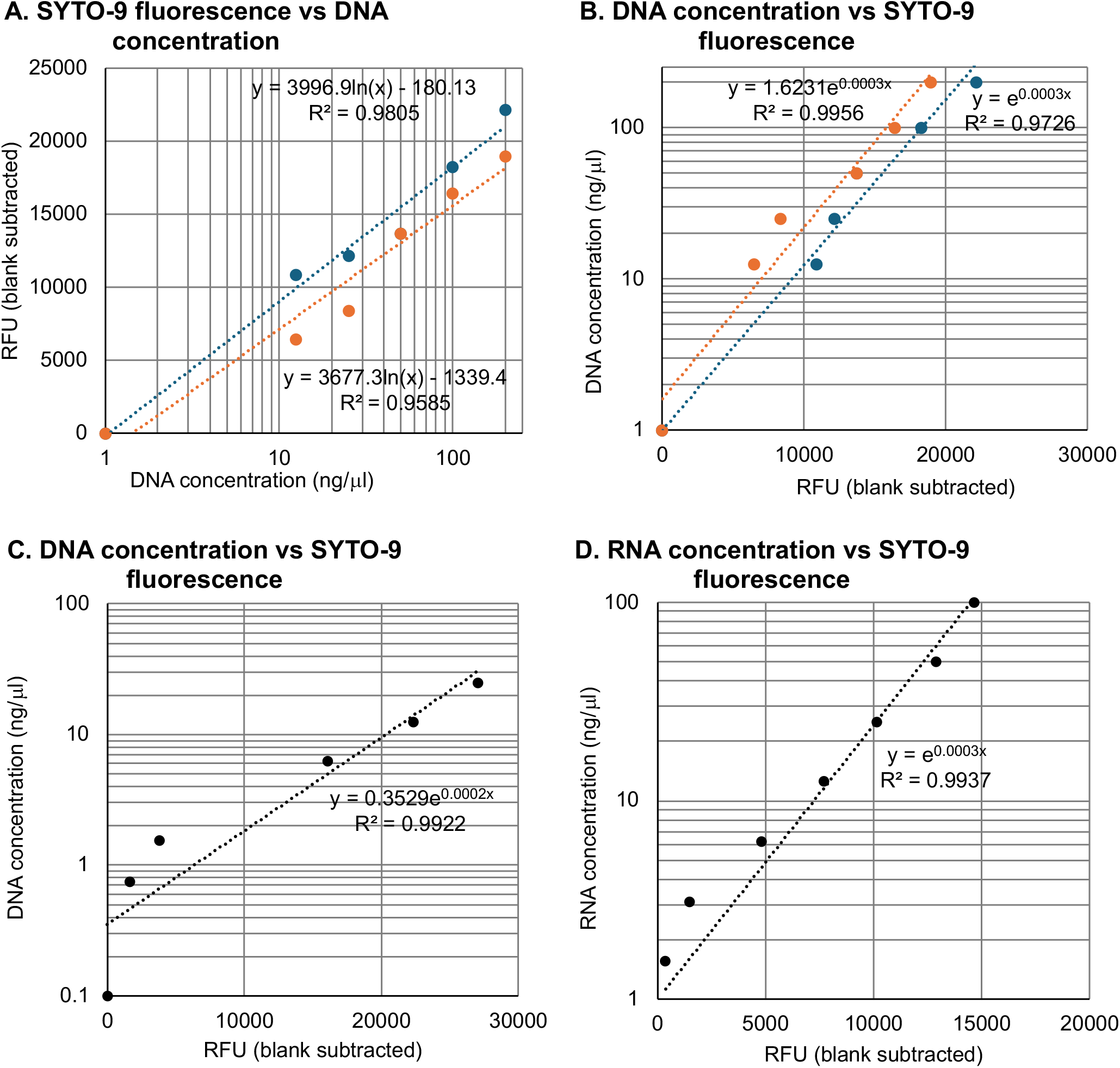
Correlations between SYTO-9 fluorescence levels and serial dilutions of nucleic acid standards. Concentration is plotted on a log scale, blank-subtracted relative fluorescence (RFU) plotted on a linear scale. Trendlines with equations and regression R^2^ values are presented for each series. **A**: Dependence of RFU on concentration for a representative series of DNA standard dilutions (12.5, 25, 50, 100 and 200 ng/µl), using a 2.5 µM SYTO-9 working stock, first read (blue markers) and 7th read (orange markers). **B**: The inverse relationship (dependence of concentration on RFU) for the series in A. **C**: Dependence of concentration on RFU for a representative series of DNA standard dilutions (0.75, 1.56, 6.25, 12.5 and 25 ng/µl), using a 10 µM SYTO-9 working stock. **D**: Dependence of concentration on RFU for a representative series of RNA standard dilutions (1.56, 3.12, 6.25, 12.5, 25, 50 and 100 ng/µl), using a 5 µM SYTO-9 working stock.

These measurements were highly stable in the face of changes to incubation times, temperatures of up to 95°C, and repeated measurements to induce photobleaching. Thus, the two series of measurements presented in Fig.1A-B underwent one or seven measurements, respectively, yielding similar values and ratios between DNA concentration and SYTO-9 fluorescence.

To extend the range of measurements to lower DNA concentration, a 4X more concentrated SYTO-9 working stock (10 µM) was used on serial dilutions of a 25 ng/µl DNA standard (0.75, 1.56, 6.25, 12.5 and 25 ng/µl), with the same protocol. A consistent exponential dependence of the form [DNA]=Ae^0.0002F^ was observed, with A=0.353 and R^2^ > 0.99 (Fig. 1C).

To check if the assay can also be used to quantify RNA, a 5 µM working stock of SYTO-9 was used on serial dilutions of a 100 ng/µl RNA standard (1.56, 3.12, 6.25, 12.5, 25, 50 and 100 ng/µl), with the same protocol. Here, too, a consistent exponential dependence of the form [RNA]=Ae^0.0003F^ was observed, with A=1 and R^2^ > 0.99 (Fig. 1D).

## Discussion

Evaluating the performance, limitations, and practical considerations of using qPCR instruments as surrogate plate readers for end-point fluorescence measurements is of significant methodological interest. Particular attention must be paid to dye selection, spectral compatibility, instrument calibration, and potential sources of variability, including fluorescence quenching effects (Eischeid 2011). By systematically addressing these factors, it may be possible to extend the utility of existing qPCR infrastructure beyond amplification-based assays, to routine end-point nucleic acid quantification.

Besides the ability to utilize existing devices for an additional common task, many qPCR instruments have a major throughput advantage compared to typical fluorometers. Thus, 384-well qPCR platforms (as used in this study) make it possible to simultaneously measure up to 384 samples (including standards and technical multiplicates); further scale-up is possible, e.g. 1536-well systems such as the Roche LightCycler® 1536. When coupled with automated plate loading, this method could present an attractive option for high-throughput projects requiring DNA quantification.

One example for such an application comes from aquaculture. Thus, in 2021 we have developed a qPCR assay for rapid, high-throughput genetic sexing of Russian sturgeon fry for the caviar industry, based on the AllWSex2 PCR marker described by Kuhl et al. (Kuhl 2021; Curzon 2022). The commercial workflow, which involves processing thousands of samples within a few days, relies on a compact field laboratory setup centered around a 384-well qPCR device; the ability to quantify the isolated DNA samples in the same high throughput and using the same device as the main assay is a valuable QC option for this and other high-throughput DNA assays.

More generally, the described approach could be adapted to additional fluorometric assays beyond nucleic acids.

### Study limitations

The method presented does not differentiate between DNA and RNA, therefore a “dirty” sample is expected to produce a combined signal resulting from different types of nucleic acids.

Assay performance in buffered or contaminated matrices (e.g., salts, EDTA, detergents, residual ethanol) was not evaluated and may alter fluorescence responses (Eischeid 2011; Zipper 2003).

The method was tested on two qPCR instruments from the same manufacturer (Bio-Rad) with an equivalent detection module (CFX-384). While the SYTO-9–SYBR spectral overlap suggests broader applicability, cross-platform validation across different optical configurations was not performed in this study.

## Acknowledgements

This publication is based upon work from COST Action BIOAQUA, CA22160, supported by COST (European Cooperation in Science and Technology).

